# Membrane capacitance recordings resolve dynamics and complexity of receptor-mediated endocytosis in Wnt signalling

**DOI:** 10.1101/417733

**Authors:** Vera Bandmann, Ann Schirin Mirsanaye, Johanna Schäfer, Gerhard Thiel, Thomas Holstein, Melanie Mikosch-Wersching

## Abstract

Receptor-mediated endocytosis is an essential process in signaling pathways for an activation of intracellular signaling cascades. One example is the Wnt signaling pathway, which seems to depend on endocytosis of the ligand-receptor complex for initiation of Wnt signal transduction. So far, the role of different endocytic pathways in Wnt signaling, the molecular players and the kinetics of this process are unclear. Here, we monitor endocytosis in Wnt3a and Wnt5a mediated signaling by membrane capacitance recordings of HEK293 cells. Our measurements revealed a fast and substantial increase in the number of endocytic vesicles. This endocytotic activity is specifically elicited by extracellular Wnt ligands; it starts immediately upon ligand binding and ceases over a period of ten minutes. By using specific inhibitors, we can dissect Wnt induced endocytosis into two independent pathways, where canonical Wnt3a is taken up mainly by clathrin-independent endocytosis and Wnt5a exclusively by clathrin-mediated endocytosis.

## Introduction

### Wnt pathway

Wnt signaling is an evolutionary highly conserved signaling pathway with important functions in embryogenesis, stem cell biology and many diseases, including cancer. After three decades of intensive research we understand many of the fundamental components of Wnt signaling pathways but it is still puzzling how so many different processes are regulated by only one system.

One level of complexity in Wnt signaling is provided by the network of a wide variety of ligands, receptors, co-receptors, antagonists, agonists and intracellular factors, which are deeply embedded in metazoan genomes (Niehrs, 2012; Holstein, 2012; Clevers et al. 2012; Nusse et al. 2017) and interact in a distinct manner. The secreted Wnt proteins activate different downstream pathways, which are traditionally classified as canonical (ß-catenin dependent) and noncanonical (ß-catenin independent). In the canonical Wnt pathway binding of the extracellular Wnt ligand to the Frizzled receptor leads to the formation of a complex with the co-receptor Lrp5/6 that recruits the scaffolding protein Disheveled and Axin, as well as GSK3 and several other intracellular components, to build up the Lrp6-signalosome. This inhibits the phosphorylation of ß-catenin, which normally marks it for proteasomal degradation and results in stabilization of ß-catenin in the cytosol, regulating various Wnt target genes upon translocation into the nucleus (Bilic et al., 2007).

The noncanonical Wnt/PCP pathway utilizes Frizzled receptors to activate Disheveled and regulates various downstream effectors like small GTPase’s of the Rho and Rac subfamily, the CaMKII and the PKC pathway (Veeman et al., 2003).

A further layer of complexity in Wnt signaling are cellular mechanisms, like endocytosis of the activated receptors (Metcalfe and Bienz, 2011; Feng and Gao, 2014). Wnt signaling can be inhibited when endocytosis of the ligand-receptor complex is blocked (Blitzer and Nusse, 2006; Rives et al., 2006; Seto and Bellen, 2006; Yamamoto et al., 2006 and 2008; Taelman et al., 2010; Hagemann et al., 2014; Gagliardi et al., 2014). Therefore, endocytosis is not only necessary for the degradation of ligand-receptor complexes but it is also essential for downstream signal transduction. A bulk of data have so far highlighted distinct differences in the endocytosis of canonical and noncanonical Wnt signaling. Clathrin-mediated endocytosis as well as caveolin-dependent endocytosis are involved in canonical Wnt signaling (Kikuchi et al. 2009). After formation of Lrp6-signalosomes in response to Wnt3a, these are internalized through a caveolin-mediated route (Sakane et al., 2010; Yamamoto et al., 2006; Vinyoles et al., 2014, Liu et al., 2014) but Wnt3a has also been shown to trigger clathrin-mediated endocytosis (Blitzer and Nusse, 2006; Seto and Bellen, 2006; Bryja et al., 2007; Jiang et al., 2012). Endocytosis of the canonical Wnt8 induced Lrp6-signalosome via clathrin-mediated endocytosis has been discovered in zebra fish (Hagemann et al., 2014).

For the noncanonical Wnt5a only clathrin-mediated endocytosis has been reported (Chen et al., 2003; Kim et al., 2008; Gorden et al., 2012; Kim & Han, 2007; Yu et al., 2007). Ohkawara et al. (2011) demonstrated that clathrin-dependent endocytosis of Wnt5a is essential for Wnt/PCP-signaling in Xenopus.

Knowledge of receptor-mediated endocytosis following Wnt stimulation is exclusively based on microscopic and/or biochemical analyses. These approaches only allow an investigation of the endpoint of endocytosis: They provide no information on their dynamics or on the temporal resolution of the endocytic process. An alternative method with a high spatial and temporal resolution sufficient for analyzing individual vesicle fission from the plasma membrane in living cells is provided by measurements of the membrane capacitance (Neher, & Marty 1982; Lindau & Neher, 1988). This method takes advantage of the fact that exo- and endocytosis are associated with changes in plasma membrane area, which in turn generate proportional changes in the electrical membrane capacitance (Cm). In this way, even the fusion and fission of single exo- and endocytic vesicles can be resolved in real time in single living cells (Rituper et al., 2013). Here we employ this technique for resolving Wnt receptor-mediated endocytosis with high temporal and spatial resolution in real time. Our data show for the first time on the cellular and physiological level that challenging of HEK293 cells with a canonical (Wnt3a) and a noncanonical ligand (Wnt5a) triggers an immediate increase in endocytosis of small vesicles and that both ligands use separate endocytic pathways; while Wnt5a is endocytosed by clathrin-coated vesicles, Wnt3a takes a clathrin-independent endocytic pathway.

## Results

### Characterization of receptor-mediated endocytosis of Wnt3a and Wnt5a in whole cell mode in HEK293 cells

We used recombinant human canonical Wnt3a and noncanonical Wnt5a to analyze their endocytic uptake into HEK293 cells. These cells contain the Wnt signaling machinery endogenously and exhibit a robust Wnt response (Sakane et al., 2010; Koo et al., 2012; Song et al., 2014). In a first approach HEK293 cells were incubated with an endocytosis marker, the styryl dye FM 4-64 (10 µM) and subsequently challenged with either recombinant Wnt3a or Wnt5a (5 ng/ml). The confocal data in Fig.1 A show that unstimulated HEK293 cells exhibited a constant rate of endocytosis. Addition of Wnt3a or Wnt5a to the bath strongly increased endocytosis. While the response to Wnt3a was immediate, Wnt5a stimulated endocytosis only after a lag of more than five minutes. After 30 minutes, it even exceeded the amount of endocytosis in Wnt3a treated cells (Fig. 1 B).

**Figure 1:**
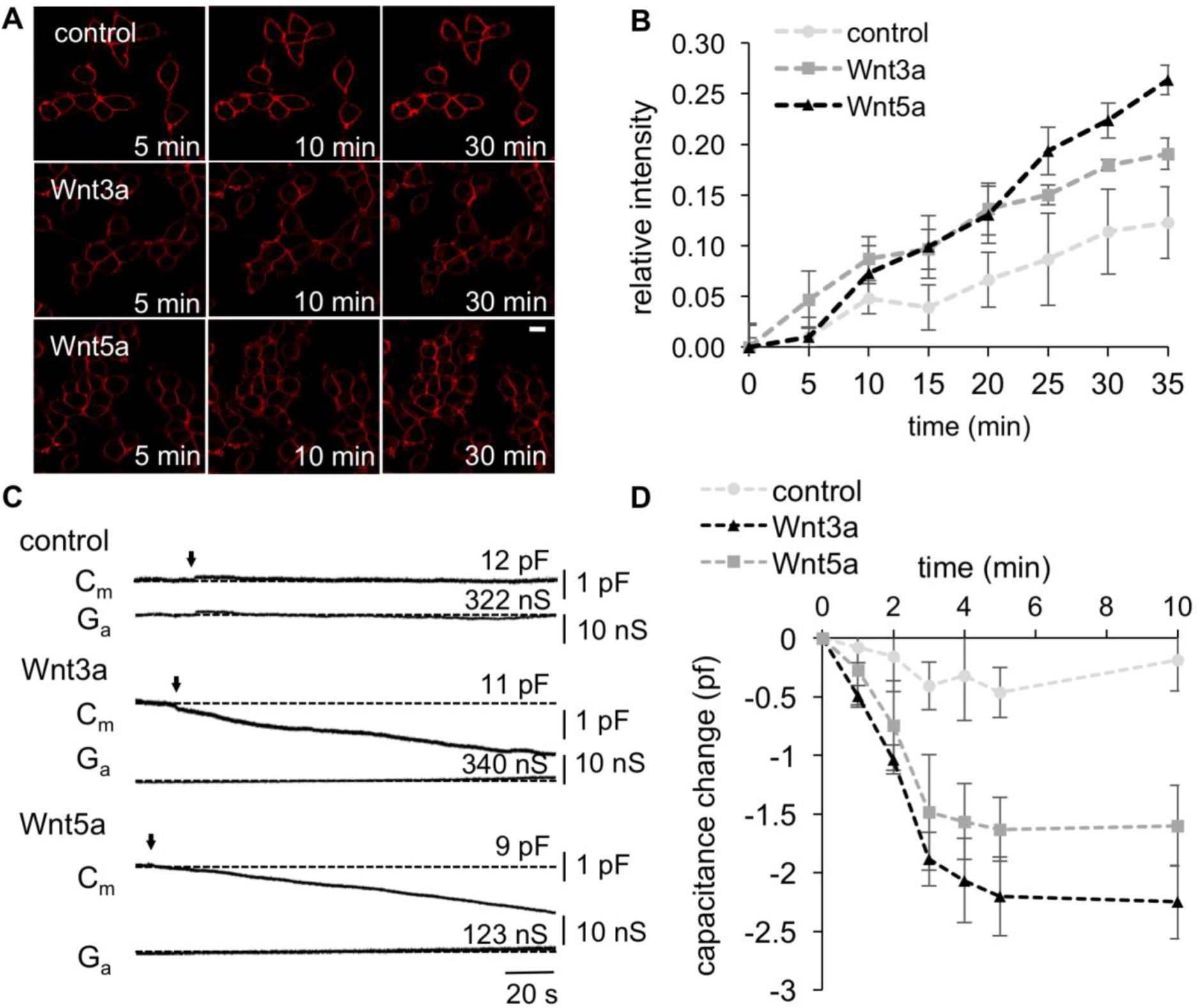
(A) Fluorescent image of FM 4-64 endocytosis in control or Wnt treated HEK293 cells. Cells were incubated for 1 minute with 10 µM of the FM 4-64 dye before addition of 5 ng/ml of the Wnt3a or Wnt5a ligands. Scale bar = 10 µm (B) Quantification of the effect of canonical Wnt3a and noncanonical Wnt5a on the uptake of FM4-64. The relative fluorescent intensity is given as the ratio of intracellular fluorescence to whole-cell fluorescence. In control cell the relative fluorescent intensity increase over time shows the steady state uptake of FM4-64. The effect of both Wnt ligands on the uptake of FM4-64 was significantly different to the control according to Student’s t-test (***, P < 0.005). Number of cells: control (n=20), Wnt3a (n=19) Wnt5a (n=18). (C) Representative whole-cell capacitance recordings of control and Wnt treated HEK293 cells. Arrow marks the time point of addition of the Wnt protein. Ga and Cm: imaginary and real part of admittance, corresponding to changes in membrane conductance (Ga lower trace) and capacitance (Cm upper trace). (D) Quantification of the effect of the Wnt ligands in whole-cell capacitance measurements. Number of measured cells: control (n=9), Wnt3a (n=9) Wnt5a (n=10).

Next, we analyzed the rapid effect of Wnt ligands on endocytosis by whole-cell patch-clamp capacitance measurements. Figure 1 C shows a representative recording of an unstimulated HEK293 cell. These cells typically exhibited a stable capacitance of 4-15 pF corresponding to a membrane area of 12-23 µm^2^. The results of these recordings show that endocytosis, which was visualized by the FM 4-64 dye in Fig. 1A, was balanced by exocytosis thereby maintaining a constant cell surface. The representative data in Figure 1 C show that treatment with Wnt3a and Wnt5a (5 ng/ml) caused an immediate and continuous decrease in the Cm value, indicating that the Wnt ligand stimulates endocytosis. In all tested cells, the Cm value dropped within one minute of stimulation by about 0.5 pF (±0.29 pF); this corresponds to a decrease in the membrane area by 4.2 µm^2^(±3.2 µm^2^). Figure 1 D summarizes the mean time course of ligand stimulated endocytosis over ten minutes. Both ligands have triggered a decrease in capacitance significantly below that of the background signal in control cells. The data further show that the ligand stimulated increase in endocytosis approached saturation already about three minutes after stimulation. Furthermore it can be noted, that Wnt3a induced a maximal drop in the Cm values (2.2 pF ±0.63 pF) that was larger than that induced by Wnt5a (1.6 pF ±0.76 pF). (Fig. 1D andere x Achse)

### Characterization of steady state exo- and endocytosis in HEK293 cells

To analyze Wnt-stimulated endocytosis at the level of individual vesicles, we performed cell-attached patch-clamp capacitance measurements. In unstimulated cells we occasionally observed spontaneous up- and downward steps of Cm, which reflect exo- and endocytosis of single vesicles respectively (Fig. 2 A-D). This type of spontaneous activity was observed in 56% of all patches (46 of 82 cells). In over 15 h of total recording time (ca. 11 min per patch) we observed on average only 2 events per cell. These spontaneous endo- and exocytic vesicles had diameters between 70 nm and 550 nm with a median of 182 nm ±149 nm.

**Figure 2:**
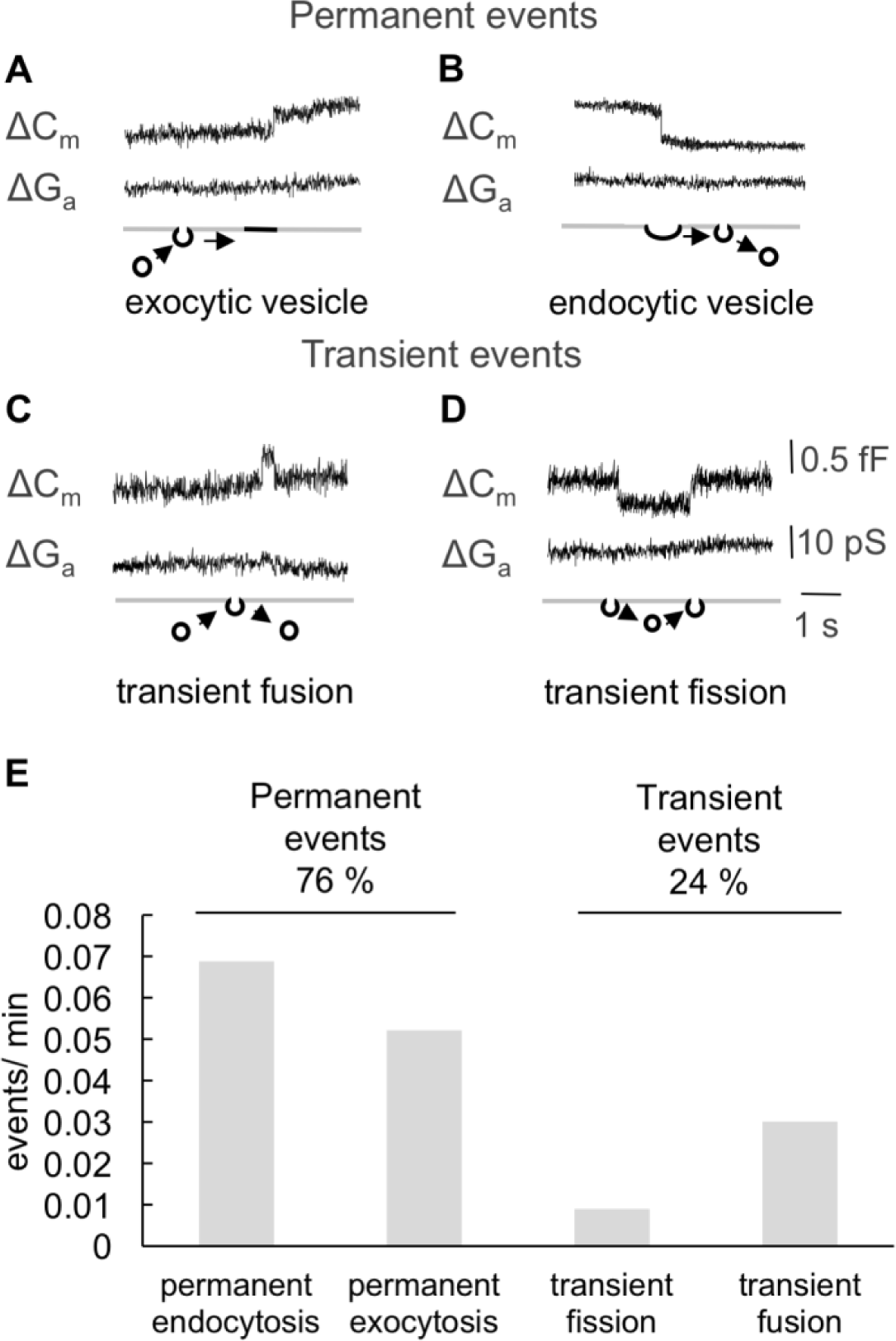
(A-D) Single spontaneous events (i.e. exocytosis, endocytosis and transients) in control HEK293 cells (imaginary and real part of admittance, corresponding to changes in membrane conductance (ΔG lower trace) and capacitance (ΔC upper trace)). (E) Frequency of permanent and transient events, number of measured cells (n=82), 15 hours recording time in total.

Spontaneous C_m_ steps could be separated according to their kinetics into four different categories (Fig. 2 A-D): permanent exocytosis (A) and endocytosis (B) and transient fusion (C) and transient fission (D). Only 24 % of the exo- and endocytic events are transient e.g. the result of so called “kiss-and-run” exo- and endocytosis (Fig. 2 E). The transient fusion of exocytic vesicles is more abundant (19 %) than transient endocytosis (5 %).

### Characterization of receptor mediated endocytosis of Wnt-treated cells

The presence of Wnt proteins in the extracellular solution (here 5 ng/ml in the pipette solution) caused a dramatic stimulation of endocytic activity (Fig 3 A). After sealing of the micropipette with the plasma membrane we monitored a robust and strong increase in permanent endocytic over exocytic activity (Fig. 3 A-C). The number of permanent endocytic events increased in this time window 17-fold for Wnt3a and 18-fold for Wnt5a over that in control cells (Fig. 3 B). This stimulation was indeed triggered only by active Wnt ligands because the inactive Wnt ligands failed to stimulate endocytosis after boiling for 60 minutes (Fig. 3B).

**Figure 3:**
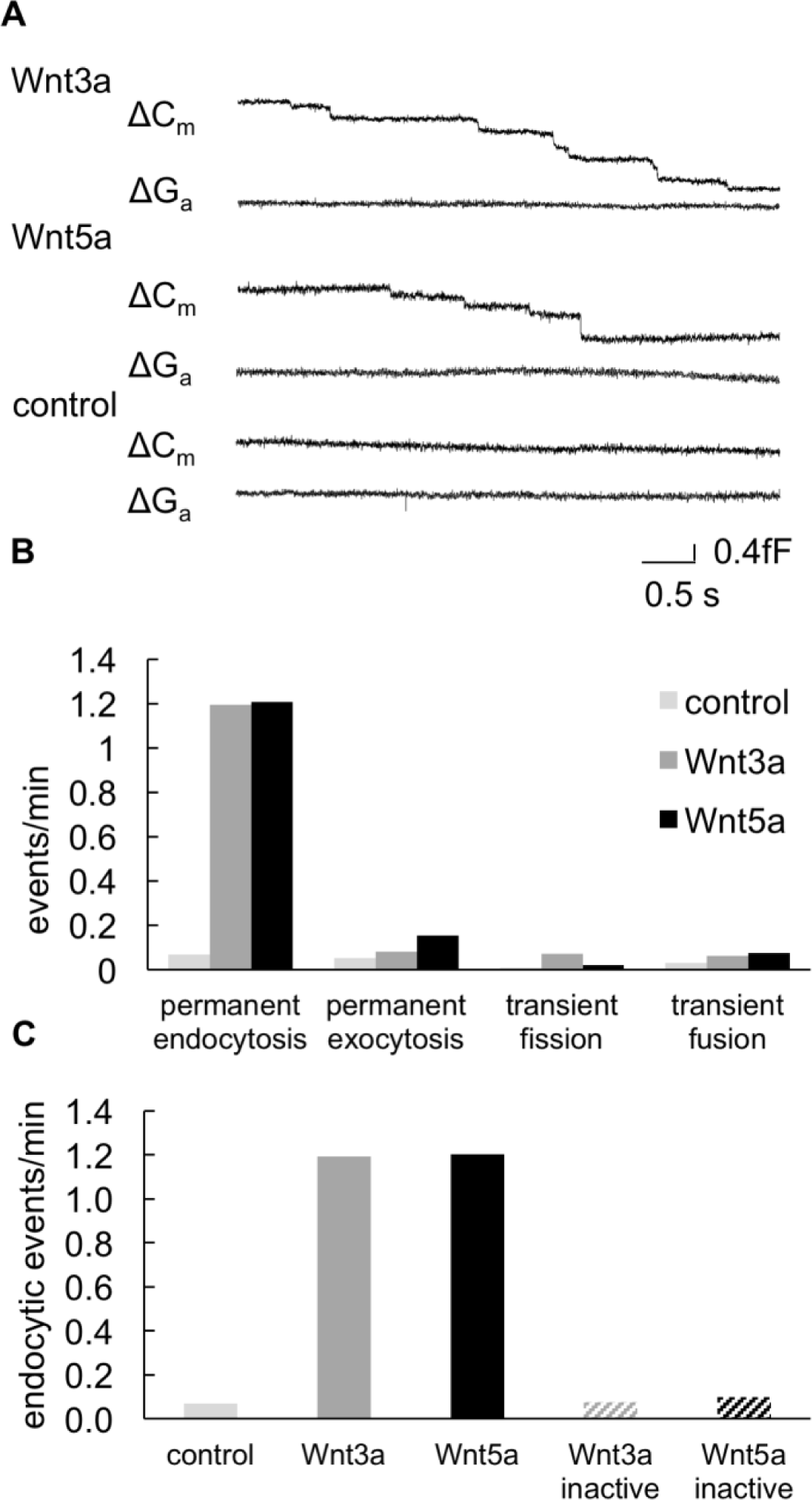
(A) Enhanced rate of endocytosis in Wnt treated HEK293 cells. Representative capacitance recordings of Wnt3a and Wnt5a treated HEK293 cells in the cell-attached mode showing successive downward steps corresponding to permanent endocytosis. Imaginary and real part of admittance, corresponding to changes in membrane conductance (ΔG upper trace) and capacitance (ΔC lower trace) for control, Wnt3a treated and Wnt5 treated HEK293 cells. (B) Number of permanent endocytic events per minute recording time in control, Wnt3a and Wnt5a treated HEK293 cells and boiled Wnt proteins. (C) Number of endocytic events per minute recording time in control, Wnt3a and Wnt5a treated HEK293 cells.

In all patches with Wnt3a or Wnt5a in the pipette solution we could observe a stimulation of permanent endocytic events. Their relative frequency increased to 85 % with Wnt3a and 83 % with Wnt5a compared to 43 % under control conditions. The degree of stimulation of endocytosis by both Wnt ligands was similar between different patches.

### Different kinetics of receptor mediated endocytosis in Wnt3 and Wnt5 treated cells

We analyzed the size distribution of endocytic vesicles, which are triggered by either Wnt3a or by Wnt5a ligands. We found that after stimulation with Wnt3a permanent endocytic vesicles had a median diameter of 109 nm (±22 nm). For Wnt5a, we observed permanent endocytic vesicles with a median diameter of 122 nm (±29 nm) (Figure 4 A). These diameters were significantly smaller than those measured in control HEK293 cells (Median 145 nm ±43 nm). This suggests that Wnt ligands were endocytosed by vesicles that were completely different from the endocytic background activity in HEK293 cells.

**Figure 4.**
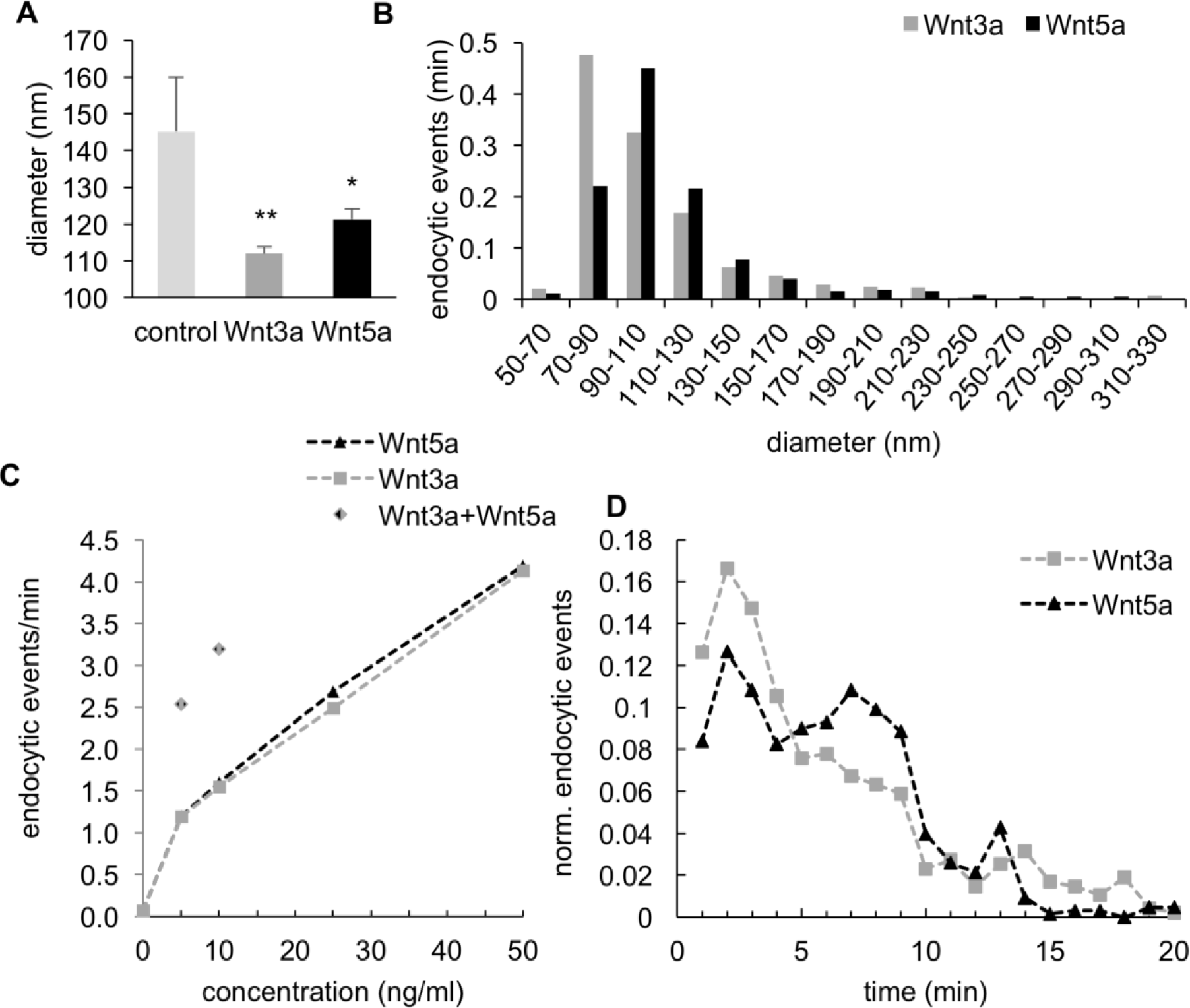
(A) Wnt-dependent size of endocytic vesicles. Endocytic vesicles stimulated by Wnt3a and Wnt5a are significantly smaller in vesicle size from the vesicles recorded in control cells (Error bars correspond to mean SEM. Asterisks indicate significant differences compared to control: ** P < 0.002, * P < 0.02; Student’s t test). (B) Distribution of different vesicle size intervals per minute recording time for both Wnt proteins. (C) Dose dependent increase of endocytic events upon addition of different concentrations of Wnt3a and Wnt5a and application of Wnt3a and Wnt5a together. (D) Temporal resolution of permanent endocytic events over 20 minutes.

To judge whether Wnt3a and Wnt5a are taken up by the same vesicle type we compared the profiles of the vesicle sizes for both ligands. This can provide a hint on the uptake mechanism since clathrin coated vesicles are generally bigger (120 nm) than vesicles in clathrin-independent endocytosis. Figures 4B shows the size distribution of Wnt3a and Wnt5a induced vesicles and indicates that Wnt3a induced vesicles are smaller in size than Wnt5a triggered vesicles.

Next, we analyzed the concentration dependency of endocytosis on both Wnt ligands. The two ligands stimulated endocytosis in a concentration dependent manner (Fig. 4C). Both Wnt’s also exhibited an additive effect when they were added together. Full saturation could not be achieved in these experiments, because ligand concentrations ≥ 50 ng/ml destabilized the seal resistance. To test whether the canonical and noncanonical ligands take the same endocytic pathway both Wnt proteins were administered together via the pipette solution at 5 and 10 ng/ml. The data in Figure 4 C show that the stimulating effect of both ligands on endocytosis is at the two concentrations tested approximately additive. The results of these experiments are best interpreted by a system in which both Wnt ligands use different and parallel endocytic pathways.

The dynamics of vesicle formation are also different for both Wnt ligands. Figure 4 D shows the temporal resolution of Wnt-induced endocytosis. Wnt3a showed a rapid increase in endocytic activity, which peaked after 2-3 minutes and declined slowly thereafter. For Wnt5a we observed biphasic kinetics of endocytosis with a first peak at 2-3 minutes and a second peak occurring after 6-8 minutes. This complex kinetic could be due to a lack of components of the endocytic machinery or the Wnt receptor complex, which have to be recycled to the plasma membrane.

Because it is well established that cells can simultaneously use different endocytic pathways, we tested to what extent independent mechanisms are acting in the endocytosis of Wnt3a and Wnt5a. To discriminate between different pathways (i.e. clathrin, caveolin- or flotillin-dependent endocytosis), we analyzed Wnt 3a and Wnt5a signaling under the influence of specific inhibitors of clathrin-dependent and clathrin-independent endocytosis. The data in Figure 5 A show that Wnt3a and Wnt5a triggered endocytosis could indeed be pharmacologically separated. Wnt3a induced endocytosis was only slightly lowered by inhibitors of clathrin-mediated endocytosis. Monodansylcadaverin and Chlorpromazin reduced endocytosis of the Wnt3a ligand by less than 10%. However, Wnt3a induced endocytosis was strongly reduced when inhibitors were used that block clathrin-independent endocytosis, e.g. Genistein and Nystatin. Both, Genistein and Nystatin reduced Wnt3a triggered endocytosis to 35.4 % or 26.3 % of the initial activity, respectively. An inverse sensitivity is found for the endocytosis of the Wnt5a protein. The endocytic activity, which is triggered by this ligand, was largely insensitive to inhibitors of clathrin-independent endocytosis; Genistein and Nystatin reduced Wnt5a triggered endocytosis by 3% and 6% respectively. Inhibitors of clathrin-dependent endocytosis in contrast caused a severe inhibition of Wnt5a triggered endocytosis reducing it by 96% (Monodansylcadaverin) or 73% (Chlorprozamine). The results of these experiments underscore the view that Wnt3a and Wnt5a are taken up into cells by two distinct and independent mechanisms. While Wnt5a enters the cell presumably via clathrin-coated vesicles, Wnt3a takes an independent route and uses a clathrin-independent endocytic pathway.

**Figure 5.**
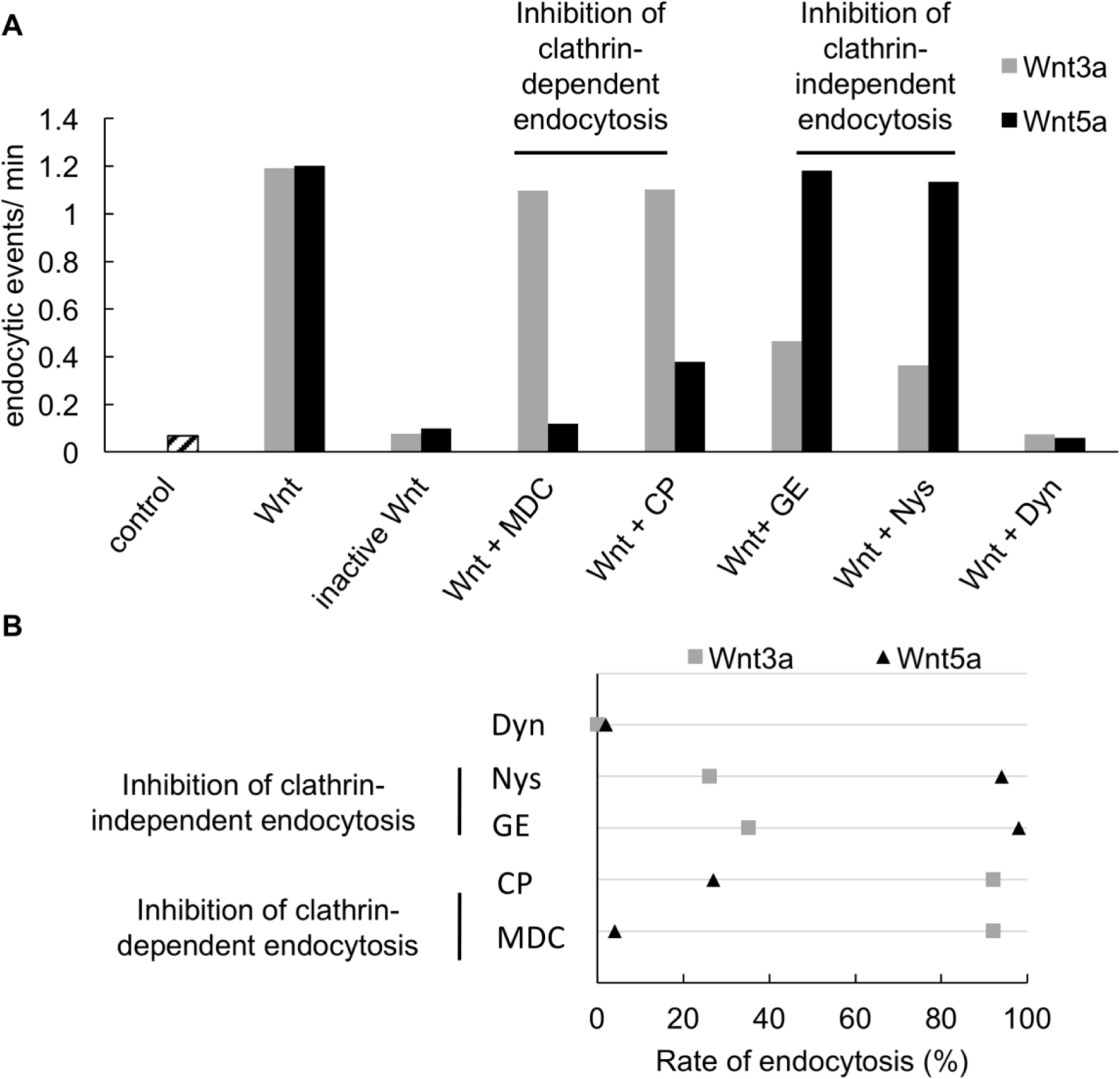
(A) Inhibition of Wnt-induced endocytosis by inhibitors of clathrin-dependent (Monodansylcadaverin MDC and Chlorprozamine CP) and clathrin-independent endocytosis (Genistein GE and Nystatin Nys). Number of permanent endocytic events per minute in control, Wnt3a and Wnt5a treated HEK293 cells with the different inhibitors. Wnt3a cannot be blocked by MDC and CP, inhibitors of clathrin-dependent endocytosis, but is blocked by GE and Nys, inhibitors of clathrin-independent endocytosis. Wnt5a-induced endocytosis can only be blocked by MDC and CP, inhibitors of clathrin-dependent endocytosis. Dynasore (Dyn) blocked Wnt3 and Wnt5a endocytosis (B) Rate of inhibition of Wnt3a and Wnt5a receptor-mediated endocytosis

To test whether endocytosis is dependent on dynamin, a small GTPase involved in scission of newly formed vesicles at the plasma membrane, we used the dynamin inhibitor Dynasore. Dynasore blocks both Wnt3a and Wnt5a induced receptor-mediated endocytosis completely (Fig. 5 A). It is well known that Dynamin is essential for clathrin-mediated endocytosis but not all forms of clathrin-independent endocytosis use dynamin. Since the receptor-mediated endocytosis of canonical Wnt3a is also completely blocked by Dynasore, dynamin seems to be essential for both endocytic routes.

## Discussion

This study shows direct *in vivo* evidence for receptor-mediated endocytosis in Wnt signaling. High-resolution capacitance measurements report an enhanced fission of single endocytic vesicles in real time which is triggered in a specific manner by relevant Wnt ligands. The frequency of permanent endocytosis events is enhanced roughly 18-fold over the background activity by the canonical Wnt3a and the noncanonical Wnt5a proteins in untreated HEK293 cells. These results demonstrate that Wnt signaling induces a fast and massive increase in permanent endocytotic activity. The fact that transient endocytotic activity was not affected to the same extend suggests that Wnt receptor-mediated endocytosis relies mostly on a permanent uptake of vesicles from the plasma membrane. A quantification of the size of the endocytic vesicles, which where induced by the Wnt ligand, shows that they are significantly smaller than endocytotic vesicles recorded under control conditions. This suggests that the Wnt ligand is not increasing the fission frequency of vesicles, that are endocytosed in a constitutive manner. The distinct size of vesicles, which are endocytosed in the presence of the ligand rather suggests that a specific type of vesicles is recruited by a highly-regulated receptor-mediated endocytosis.

The Wnt triggered mechanism of receptor-mediated endocytosis can be further dissected into two discrete routes involving or not involving clathrin with pharmacological experiments using specific inhibitors. The uptake of Wnt5a strongly depends on clathrin-mediated endocytosis, while Wnt3a is internalized to a great extent via clathrin-independent mechanisms. The receptor-mediated endocytosis of Wnt3a could be blocked to a great extent with Genistein and Nystatin both inhibitors of clathrin-independent endocytosis. This is in accordance with several studies which showed that canonical Wnt3a signal activation is clathrin-independent (Sakane et al. 2010, Liu et al 2014, Vinyoles et al. 2014). Inhibition of clathrin-mediated endocytosis by Monodansylcadaverin and Chlorprozamine on the other hand leads to a strong reduction of receptor-dependent endocytosis of Wnt5a. It has been assumed that blocking of one pathway of receptor-mediated endocytosis may promote entry through another pathway, which is not important under physiologically conditions (Pelkmans et al. 2005). The present data however show that Wnt5a is not endocytosed by any alternative endocytic pathway when clathrin-mediated endocytosis is inhibited. This underscores that Wnt5a enters the cell exclusively via clathrin coated vesicles. For Wnt3a the situation is more complex. Blocking of clathrin-dependent endocytosis caused a small 10% reduction in endocytosis. This suggests that Wnt3a can enter cells to a small extent also via clathrin-mediated endocytosis. Several studies indeed suggest also a role of such clathrin-mediated endocytosis in canonical Wnt3a signaling. Yamamoto et al. (2008) showed that clathrin-mediated endocytosis is important for Wnt3a signal inhibition by sequestration of its receptor LRP5/6 from the plasma membrane in response to the Wnt antagonist Dkk1.

Dynasore blocks the uptake of both Wnt proteins which underlines the assumption that Dynamin plays a key role in canonical and noncanonical Wnt receptor-mediated endocytosis, independent of the vesicular coat proteins or the endocytic pathway. Dynamin is known to be important in clathrin and caveolin dependent endocytosis (Viera et al., 1996).

The concept of two parallel and mutually independent pathways for endocytosis of canonical and noncanonical Wnt signaling is further supported by experiments in which both ligands were administered together. Each of the Wnt proteins induces by itself endocytosis in a concentration dependent manner. The application of both Wnt proteins together shows an additive effect in the sense that the total frequency of endocytosis exceeds the maximal frequency, which is achieved by an individual ligand. The evidence for parallel and mutually independent pathways is further underscored by our finding that specific inhibitors of one pathway do not interfere with the other endocytotic pathway. Both Wnt proteins do not depend on the same pathway for endocytosis because the number of endocytic vesicles is not increasing linearly with the ligand concentration. Instead, the frequency of endocytosis reaches a saturation level, which can be only overcome by adding both Wnt ligands together. This behavior would not be expected if both Wnt ligands were preferentially interacting with the same pathway. Our finding of two independent endocytotic pathways is different from previous hypotheses postulating that the suppression of Wnt5a by Wnt3a can be explained by the competition of both ligands for the same Frizzled 2 receptor at the plasma membrane (Sato et al, 2010).

Receptor-mediated endocytosis, which is triggered by both Wnt ligands, is a very fast process. The temporal resolution of the ligand triggered endocytosis indicates that vesicle fission occurs immediately upon receptor binding and ceases again after a few minutes. This fast endocytosis argues for an active role of endocytosis in canonical as well as noncanonical Wnt signaling. The whole signaling complex, as well as the receptor-endocytosis machinery, must be present in close proximity to the plasma membrane. This assumption is consistent with data from Wang et al. (2016) who showed that endogenous Axin, a crucial component of the early Wnt pathway, is already present at the plasma membrane in puncta in Drosophila without stimulation through the Wnt pathway. The wave like dynamics of Wnt5a receptor-mediated endocytosis, which is seen in Fig. 4 C, indicates that a pool of essential components in the endocytotic machinery, may indeed be present in unstimulated cells but used up during the initial burst of endocytotic activity. This pool may need recruitment of new components to the plasma membrane to sustain active endocytosis.

In the case that receptor mediated endocytosis determines the onset of the Wnt signaling cascade, one might expect that downstream signaling events occur only after this event. The first step after binding of Wnt to its receptor is the activation of GSK3 which leads to the formation of the LRP6-signalosome, phosphorylation and recruitment of Axin to this complex and the ensuing stabilization of ß-catenin in the cytosol followed by its translocation into the nucleus.

Recent data on the time scale of the intracellular canonical Wnt signaling cascade indeed indicates a dynamic that is in good agreement with receptor-mediated endocytosis detected in our measurements. For canonical Wnt3a signaling Ding et al. (2000) showed a GSK3 activity within the first ten minutes after Wnt addition. The phosphorylation of Axin is detectable shortly after Wnt 3a treatment in HEK 293 cells and is diminished after 15 to 30 minutes (Kim et al., 2013; Yang et al., 2016). This means both events follow directly after the endocytic activity is observed in our measurements.

This sequence of events highlights the fact that the fast and vesicle specific endocytosis of Wnt ligands occurs as a first and initial step guiding the intracellular Wnt signaling cascade towards canonical and noncanonical Wnt signaling.

## Conclusion

In this study we show direct evidence, for receptor-mediated endocytosis in Wnt signaling. High resolution membrane capacitance measurements are able to resolve a very fast kinetics of Wnt3a and Wnt5a uptake in real time. By using specific inhibitors, we could distinguish between canonical and noncanonical pathways and demonstrate that the internalization of Wnt5a strongly depends on clathrin-mediated endocytosis while Wnt3a uptake is mostly driven by clathrin-independent mechanisms. The crucial role of dynamin in both endocytic pathways could be shown by an almost complete block in endocytosis after dynasore treatment. Since receptor mediated endocytosis plays also an important role in many other morphogen signaling pathways like Notch, Hedgehog and TGFß-signaling we expect many more applications for high resolution patch-clamp capacitance studies.

Taken together, these results underline the importance of receptor-mediated endocytosis in Wnt signaling and provide the basis for identifying new molecular players in the early differentiation of canonical and noncanonical Wnt signaling.

## Materials & Methods

### Electrophysiology

Patch pipettes with a tip resistance in the range of 5 MΩ were prepared daily from glass capillaries (Kimax 51, Kimax Products, Vineland, NJ, USA), which were coated with Sigmacote (Sigma-Aldrich, Munich). Pipettes were filled with bath solution.

Measurements were performed with a dual-phase lock-in patch-clamp amplifier (SWAM IIC, Celica, Ljubljana, Slovenia). Membrane patches were clamped at 0 mV on which a sine wave (root mean square 111mV, sine wave frequency 1.6 kHz) was applied. The phase of the lock-in amplifier was adjusted to nullify changes in the real part (Re) of the admittance signal to a manually generated 100 fF calibration pulse. The output signals were low pass filtered (10 Hz, -3 db), acquired at 100 Hz by an A/D converter (NI-DAQ, National Instruments, Austin, USA) and stored on a personal computer. The signal in-phase reflects Re and is equivalent to the patch conductance; the out-of-phase signal corresponds to the imaginary part (Im). If there is no reflection in the Re trace the Im trace is directly proportional to the Cm changes. For events with projections between Re and Im the vesicle capacitance Cv and the pore conductance Gp were calculated according to Lollike and Lindau (1999):

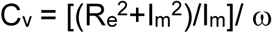

Where ω is the angular frequency (ω=2πf) and

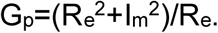

As the membrane capacitance is proportional to the membrane area, the diameter (d) of the vesicle can be determined from the vesicle capacitance according to the equation

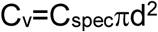

where C_spec_(specific capacitance) is the capacitance per unit membrane area. It was set to 0.8 µFcm^-2^, a value that has previously been described for cellular membranes (Gentet et al., 2000; Neher & Marty, 1982). Events were analyzed using the cursor option in the software subroutine (CellAn, Celica, Slovenia) written for MATLAB (MathWorks Inc., Natik, MA, USA) with additional digital filtering as required. Values are presented as median or mean ± SEM. Data were stored as described previously (Bandmann and Homann 2012) in a MySQL Community Server 5.1.49 database (Oracle). Calculations were performed using the CAMMC web application and results were again stored in the database.

For whole-cell measurements we only analyzed data sets with a stable access conductance G_a_ over 100 nS during the entire measurement (45 min).

Therefore, HEK293 cells were kept in an extracellular bath solution and the membrane patch of approximately 1 µm^2^ under the pipette stayed intact.

### Confocal imaging

HEK293 cells were imaged with a Leica TCS SP5 II spectral confocal microscope (Leica Microsystems). Images were acquired with an HCX PL APO CS 40×1.3 Oil UV object.

Cells were incubated in 10 µM FM 4-64 (Invitrogen, Darmstadt, Germany) for different time points and excited with the 488 nm line of a 25 mW Argon laser and fluorescence was detected at 630-700 nm. Images were analyzed with ImageJ software (NIH). For analysis of the relative intracellular fluorescence the ratio of intracellular fluorescence (area below the outline of the plasma membrane) compared with the whole cell fluorescence was used. Values are presented mean ± SEM.

### Cell culture

HEK293T cells were grown in continuous culture as previously described (Mikosch et al., 2006). Recordings were made within 1-3 days after plating. Experiments were performed on cells incubated at 37°C in 5 % CO2 for 2 to 3 days. Cells were bathed in a bath solution containing the following: 20 mM KCl, 1.8 mM CaCl2, 1mM MgCl2, and 5 mM HEPES at pH 7.4. Mannitol was used to adjust the osmolarity to 300 mOsmol/kg.

The purified recombinant Wnt3a and Wnt5a proteins were suspended with PBS, 0.1 mM EDTA, 0.5 % CHAPS and 0.5 mg BSA (R&D Systems). They were used at 5, 10, 25 and 50 ng/ml.

### Inhibitors

Clathrin-dependent endocytosis was analyzed by using Chlorprozamine hydrochloride (CP) and Monodansylcadaverin (MDC). Chlorprozamine hydrochloride (100 µM for 10 minutes) is a cationic amphiphilic drug that inhibits clathrin-mediated endocytosis by depleting clathrin and adapter protein 2 (AP2) from the plasma membrane (Qian et al., 2002; Vercauteren et al., 2010; Firdessa et al., 2014). Monodansylcadaverin (MDC) (1mM for 30 minutes) blocks the enzyme transglutaminase 2, which is necessary for receptor crosslinking in the region of clathrin-coated pits. Both, CP and MDC have been shown to selectively inhibit clathrin-mediated endocytosis (Okamoto et al., 2000).

Caveolae-mediated endocytosis was inhibited by using Nystatin, Methyl-ß-cyclodextrin and Genistein. Nystatin and Methyl-ß-cyclodextrin (MßCD) deplete the membrane of cholesterol and therefore are widely used to inhibit caveolin-mediated endocytosis. Our experiments with Nystatin (25 µg/ml for 30 minutes) and MßCD (5 mM for 30 minutes) revealed that HEK293 cells are heavily affected by both treatments. After treatment with 5 mmol/l MßCD we were not able to perform regular patch-clamp measurements because the formation of a GΩ-seal was not possible or the seal disrupted after a few minutes. The rate of transient events increased and permanent exocytosis was reduced to almost zero which fits to the idea that cholesterol plays also an important role in docking and fusing of exocytic vesicles (Rituper et al., 2012). We also tested Genistein (200 µM for 30 minutes), which is a tyrosine kinase inhibitor that causes local disruption of the actin network at the site of endocytosis and effectively inhibits caveolae-mediated endocytosis (Vercauteren et al., 2010; Dos Santos et al., 2011; Rejman et al., 2004). 200 µM Genistein did not show apparent toxic effects on HEK293 cells. Dynasore was used at 80 µM for 30 minutes and did not show toxic effects on HEK293 cells.

## Acknowledgments

We thank Ulrike Homann and Kerri Kukovetz for valuable comments on the manuscript. No conflict of interest declared.

